# Alterations in protein kinase networks in astrocytes and neurons derived from patients with familial Alzheimer’s Disease

**DOI:** 10.1101/2022.06.14.496149

**Authors:** Nicholas D. Henkel, Alex Joyce, Elizabeth Shedroff, Ali Sajid Imami, Khaled Alganem, Abdul-rizaq Hamoud, Chongchong Xu, Benjamin Siciliano, Tao Ma, Zhexing Wen, Robert E. McCullumsmith

## Abstract

Neurons and astrocytes derived from Alzheimer’s Disease (AD) patient induced pluripotent stem cells are an evolving technology used to study the pathogenesis and etiology of AD. As the utility of mouse models of AD are increasingly coming into questions, using iPSC technology may offer an opportunity to study this disease with human substrates. Herein, we using a hypothesis generating platform, the PamGene12 Kinome Array, to identify core protein kinases in neurons and astrocytes derived from familial AD patient iPSCs. We identified five core protein kinases in these cells and examined the pathways in which they are enriched. Importantly, we complement our findings using an in-silico approach with postmortem AD brain datasets. While these protein kinases have been conceptualized in the context of traditional AD pathology, they have not been explored in the context of aberrant signaling in the pathophysiology of the disease.

## Introduction

Alzheimer’s Disease (AD) is a devastating neurodegenerative disorder characterized by the gradual and insidious loss of memory, cognition, praxis, executive and physiologic function. AD is the most common cause of dementia in the United States (1). Since its characterization in 1907, it has been intensely researched.

On histopathology, AD is famously characterized by the accumulation of amyloid-beta (Aβ) and hyperphosphorylated tau (2). Though the presence of these oligomers is toxic to the brain, it does not fully explain how AD develops (3, 4). That is, Aβ nor tau alone are sufficient to induce AD: both can be present in aged brains without a dementia diagnosis (3, 5, 6). Despite this, mouse models harboring human transgenes in pathways related to amyloid and tau processing have been the pillar of AD research for the past thirty years. While the research generated from these mice have provided a wealth of information on the pathophysiology of AD, the ability of these models to either reliably recapitulate the human neurodegenerative phenotype of AD or produce efficacious therapies for AD has failed.

In the past fifteen years, the advent of induced pluripotent stem cell (iPSC) technology has changed how neurodegenerative diseases, such as AD, are studied. The differentiation of iPSCs to neural progenitor cells followed by neuronal, glial, and vascular specialization have closely modeled pathology of the human brain. Neurons and astrocytes, for instance, derived from AD-patient iPSCs exhibit elevated amyloid beta production, impaired autophagy, defective synaptic function, and oxidative stress (7–16). The technology to model iPSCs for human disorders is evolving; however, these cells provide an opportunity to expand our understanding of the pathophysiology and pathogenesis of human AD. While postmortem brain has been an invaluable tool to understand the AD brain, there is no opportunity for mechanistic or temporal evaluation and analysis. iPSCs provide an avenue to overcome this limitation.

14 million Americans are expected to develop AD by 2060 (1). Despite the acceleration of AD prevalence, there are no effective treatments, and many recent clinical trials have failed (17, 18). To identify disease-altering targets, a deeper understanding of the pathophysiology of this devastating illness is needed.

Individual elements of intracellular protein kinase signaling have been studied in AD. However, protein kinase-mediated signaling events are not linear cascades and should not be studied in isolation (19). They include myriad protein kinases, phosphatases, and signal integration molecules that form networks (together, forming the “kinome”) that coordinate cellular functions and pathological fates of cells. For this reason, understanding the AD kinome will be crucial to identify novel pathophysiological insights.

Most of the protein kinase studies in AD have either focused on kinase expression changes, the direct impact of the kinase(s) on Aβ and/or tau pathology, or interpret their data considering protein kinases as individual components of linear cascades. Further, very few AD studies have employed network level approaches to measure protein kinase expression or function. Several network-based studies have deployed mRNA datasets, proteomic studies, or have focused on specific signaling hubs, such as WNT and mTOR, to study the AD kinome (20–28). Expanding our understanding of the AD pathophysioloy will require approaching the problem agnostic to a priori hypotheses related to AD.

To overcome this, we deployed the PamGene12 Kinome Array to study the kinome in neurons and astrocytes derived from three familial AD patient iPSCs. In this study, we have identified 5 common kinases between these familial AD mutation cell lines that may represent core hubs in the disease pathophysiology. Interestingly, these kinases are enriched in pathways that mediate metabolism and metabolic functioning. Finally, we deployed several omics datasets from postmortem brain and a neuroglioma cell line to confirm and complement our findings.

## Methods

### Culture of Human iPSCs and differentiation into cortical neurons and astrocytes

Human iPSC lines were previously generated and fully characterized (29) (Figure 1). All experiments were performed in compliance with the relevant laws and institutional guidelines. Human iPSCs were differentiated into cortical neurons following the previously established protocol (29–32). Briefly, hiPSCs colonies were detached from the feeder layer with 1 mg/ml collagenase treatment for 1 hour and suspended in embryonic body (EB) medium, consisting of FGF-2-free iPSC medium supplemented with 2 μM Dorsomorphin and 2 μM A-83, in non-treated polystyrene plates for 4 days with a daily medium change. After 4 days, EB medium was replaced by neural induction medium (NPC medium) consisting of DMEM/F12, N2 supplement, NEAA, 2 μg/ml heparin and 2 μM cyclopamine. The floating EBs were then transferred to matrigel-coated 6-well plates at day 7 to form neural tube-like rosettes. The attached rosettes were kept for 15 days with NPC medium change every other day. On day 22, the rosettes were picked mechanically and transferred to low attachment plates (Corning) to form neurospheres in NPC medium containing B27. On day 24, the neurospheres were then dissociated with Accutase at 37°C for 10 minutes. For cortical neuron differentiation, dissociated neurospheres were placed onto Poly-D-Lysine/laminin-coated coverslips in the neuronal differentiation medium, consisting of Neurobasal medium supplemented with 2 mM L-glutamine, B27, cAMP (1 μM), L-Ascorbic Acid (200 ng/ml), BDNF (10 ng/ml) and GDNF (10 ng/ml). Half of the medium was replaced once a week during continuous culturing. For astrocyte differentiation, dissociated neurospheres were plated on the Matrigel coated plate in astrocytes medium (ScienCell). After the astrocytes projected outside the neurospheres and populated the plate surface, the neural progenitor spheres were pipette off the plates using 1 ml pipettes and transferred to new Matrigel coated plates for continuous generation of astrocytes. Then cells surrounding the neurospheres were dissociated enzymatically using TrypleE (Invitrogen) and plated in the 100 ug/ml PDL (Sigma)-coated cell culture dish.

**Figure 1.**
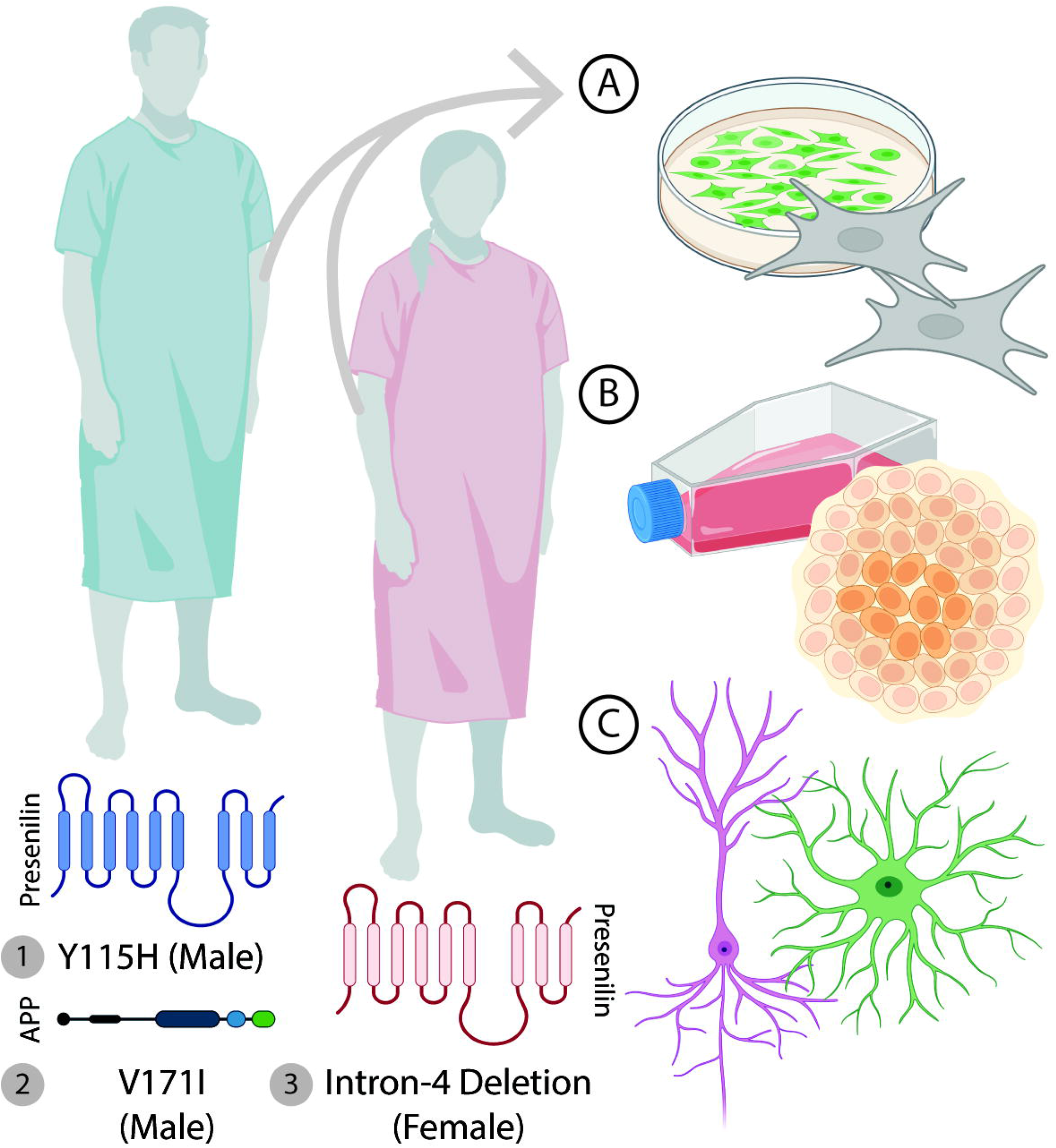
Differentiation of Alzheimer’s Disease human fibroblasts to induced pluripotent stem cells. Fibroblasts from three Alzheimer’s Disease patients were harvested for analysis. Two male cell lines (Presenilin-1 Y115H; Amyloid Precursor Protein V171I) and one female (Presenilin-1 Intron-4 deletion) were used and compared to non-isogenic controls. Differentiation of collected human fibroblasts were cultured (A), differentiated to induced pluripotent stem cells (B), then differentiated to cortical neurons and astrocytes (C).

### Sample preparation for Kinome Array profiling

Cell lysates were homogenized in a cocktail of Mammalian Protein Extraction Reagent (MPER, ThermoFisher Scientific) and 1:10 HALT Protease and Phosphatase Inhibitor (ThermoFisher Scientific). Protein concentrations were determined used the Pierce BCA Reagent Kit (ThermoFisher Scientific). Lysates were taken to 1.0 ug/uL, aliquoted, and stored at −80° until the day of the experiment. For profiling on the Serine/Threonine PamChip, samples were taken to 0.2 ug/uL and loaded onto three chips. Each chip contains 4 wells. Samples were run in singlicate.

### Kinome Array Profiling

The raw PamChip images are pre-processed using BioNavigator (PamGene International B.V.) to generate numerical values of the signal intensity for each reporter peptide on the chip. BioNavigator deploys the Upstream Kinase Analysis (UKA) tool from assign upstream kinases. The default settings of the standard STK analysis protocol was used with the additional step of upstream kinase analysis. UKA reports the final score as a metric for ranking implicated kinases, which is calculated based on the specificity of the peptides mapped to the kinases and the significance of phosphorylation changes of the peptides between the compared groups. To analyze the kinome array profiles, we further deployed the Kinome Random Sampling Analyzer (KRSA) R package to pre-process, apply quality control checks, and select differentially phosphorylated peptides (33). KRSA was used to analyze the kinome profiles of our cell lines. The analysis starts with calculating the linear regression slope (signal intensity as a function of camera exposure time) followed by scaling the signal by multiplying by 100 and then log transforming the scaled values. The derived values from the previous steps are used as the final signal (i.e., peptide phosphorylation intensity) in the comparative analyses. Additional quality control filtration steps were carried out to remove peptides with either very low signal intensity (signal < 5) or R2 of less than 0.9 in the linear model. The signal ratios between pairs of groups (control versus AD-patient derived iPSCs) were used to calculate log2 fold change (LFC) for each peptide. The LFC was calculated per chip and by sex, and then averaged across chips. To compare the kinase activity, peptides with an average LFC of at least 0.2 were carried forward to the upstream kinase analysis. To investigate associated upstream kinase families, KRSA takes the list of differentially phosphorylated peptides and uses a random resampling approach to assign scores for each kinase family.

### In-Silico Data Exploration

The Enrichr web tool was used to perform gene set enrichment analyses (34). Comparisons across genotypes and cell types were manually curated using the Bioinformatics and Evolutionary Genomic Venn Diagram tool (35). The peptides from the Kinome Array are mapped to their corresponding proteins, then to their official HGNC gene symbols. KEGG_2021_Human was used for pathways analysis. Further, the web tool Kaleidoscope was used to query multiple curated databases and the Integrative Library of Integrated Network-Based Cellular Signatures (iLINCS) (36–38). Using our kinome data from UKA, we uploaded our kinases and their scaled scores to KinMap to generate the kinome phylogenetic trees (39).

## Results

### Perturbed phosphorylation of serine/threonine kinases in neurons and astrocytes derived from AD-patient iPSCs

Cell lysates from neurons and astrocytes were on the PamGene12 Kinome Array. There are 144 peptide substrates blotted on the STK PamChip. As a quality control measure, peptides that were un-detectable or did not increase in signal linearly with exposure time in the post-wash phase were eliminated for each comparison respectively. The peptides that met inclusion criteria were carried over for analysis. Globally, there is a perturbation of phosphorylation intensity in the neurons and astrocytes derived from the AD-iPSCs (Figure 2A, Supplemental Figure 1E). We detected a significant reduction in global phosphorylation intensity in the astrocytes derived from both the Male PS1 Y115H iPSCs (Figure 2B, p = 0.035), and the Female Intron-4 deletion iPSCs (Supplemental Figure 1C; p = 0.017). However, there was no difference in the phosphorylation intensity in the astrocytes derived from the APP V717I iPSCs (Supplemental Figure 1D; p = 0.14). Further, there was no reduction in the phosphorylation intensity in the neurons derived from the neither Male PS1 Y115H iPSCs (Figure 2B, p = 0.49) nor the Male APP V717I iPSCs (Supplemental Figure 1B, p = 0.14). Only the neurons derived from Female Intron-4 deletion iPSCs displayed a significant reduction in phosphorylation intensity (Supplemental Figure 1A, p = 1.2E-5). These findings suggest both cell type and genotype specific deficits in serine/threonine protein kinase activity.

**Figure 2.**
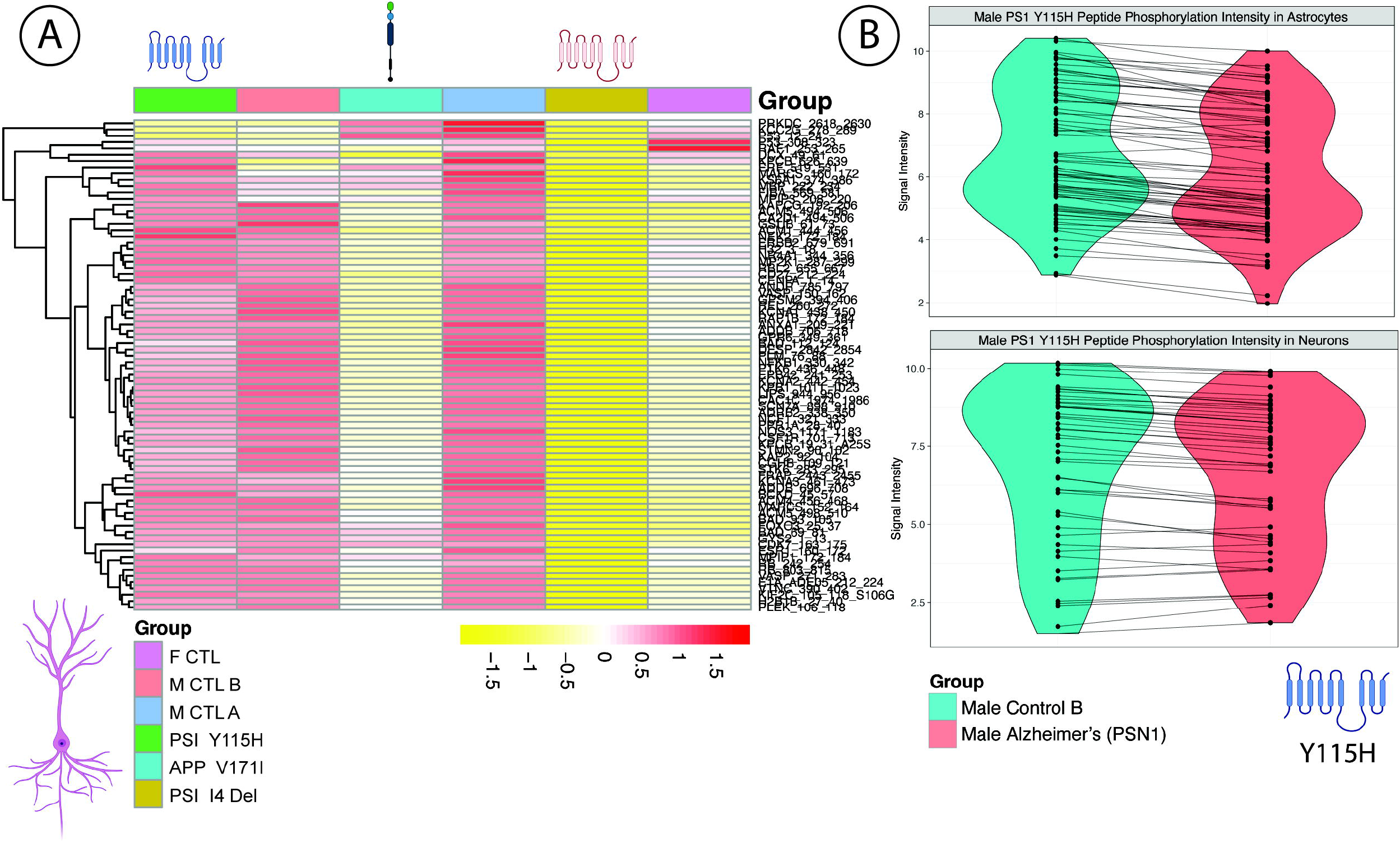
Global peptide phosphorylation is changed in neurons and astrocytes derived from AD-subject iPSCs. Representative heatmap (A) from our neurons derived from our three control and AD cell lines and violin-plots (B) of neurons (top) and astrocytes (bottom) derived from Male PSN1 Y115H iPSCs.

### Kinase assignments

Using our bioinformatic workflow, Kinome Random Sampling Analyzer (KRSA), we identified protein kinases families that are both under- and over-represented in our AD substrates. By convention of the software, we define an under-represented kinases as a kinase with a negative Z-Score (of the kinases that phosphorylate the differentially phosphorylated peptides, an under-represented kinase was observed less than the expected average). Similarly, we define an over-represented kinase as a kinase with a positive Z-Score (of the kinases that the phosphorylated peptides, an over-represented kinase was observed more than the expected average). It is crucial to note that we do not contend that an assignment of under- or overrepresentation indicates a concordant change in biological or functional activity. Rather, we purport that the KRSA assignment renders a list of kinases that may be changed in the analytical substrate, compared to the reference control, as a “hit” kinase. Defining the biological role requires traditional biochemical experiments (i.e., transcript and protein expression, as well as enzyme activity assays). However, it is reasonable to deduce that an underrepresented assignment may represent a hub that is “missing” in the disease state while an overrepresented assignment may represent a hub that remains “present” in the disease state.

To identify novel and shared kinases in our three familial AD cell lines, we compared our KRSA results from both the astrocytes and the neurons. Importantly, for our analysis, we intended to identify kinases that were similar across genotypes and cell types. Such kinases may represent core signaling hubs that are perturbed in the disease. Each disease genotype was compared to a non-isogenic control cell line from a patient with no neuropsychiatric disorder. For each comparison, we generated a complete list of under and overrepresented kinases using the peptides that met filtering parameters (Supplemental Table 1). We mapped these kinases, respectively, for each cell type and genotype on the kinome phylogenetic trees (Figure 3; Supplemental Figure 2).

**Figure 3.**
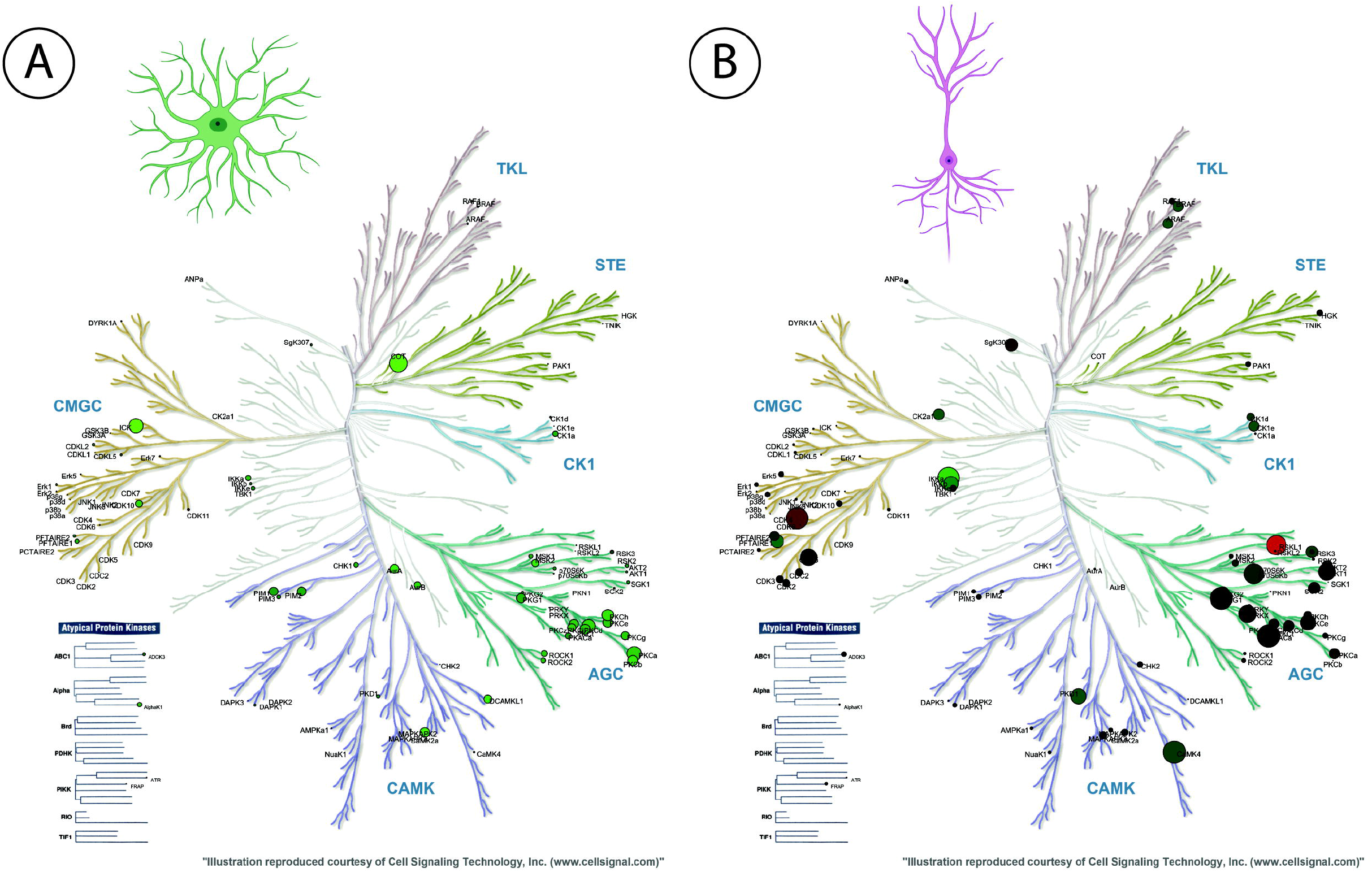
Differential dysregulation of kinase networks in AD-subject iPSCs. Representative phylogenetic trees of the “hit” protein kinases in astrocytes (A) and neurons (B) in the Male Y115H iPSCs.

### Protein kinases implicated in metabolism and metabolic function are shared across the genotypes and cell types

We examined the underrepresented and the overrepresented kinases separately to distill a list of hit kinases with greater resolution. For both the neurons and astrocytes derived from iPSCs, all three cell lines, Female PSN1 Intron-4 deletion, Male PSN1 Y115H, Male APP V717I, three protein kinase families were consistently assigned a negative Z-Score: DYRK, JNK, and PDK1 (Figure 4A). LKB, ERK, and p38 were only underrepresented in all neuron cell lines while PLK, MTOR, NMO, and GSK were underrepresented in all astrocyte cell lines (Figure 4A). While these latter 7 kinase families may have been underrepresented in a comparison cell line (control versus disease cell line), we highlight the kinases that were shared across all comparisons. On the other hand, for the overrepresented kinases, RAD53 and PIM were consistently assigned a positive Z-Score in both cell lines and all genotypes (Figure 4C). There were no unique, shared overrepresented kinases for astrocytes. DMPK, PKG, PKA, PKD, PKC, PAKA, and RSK were common overrepresented kinases in neurons only (Figure 4C). Our “intersection kinases,” DYRK, JNK, PDK1, RAD53, and PIM were further examined.

**Figure 4.**
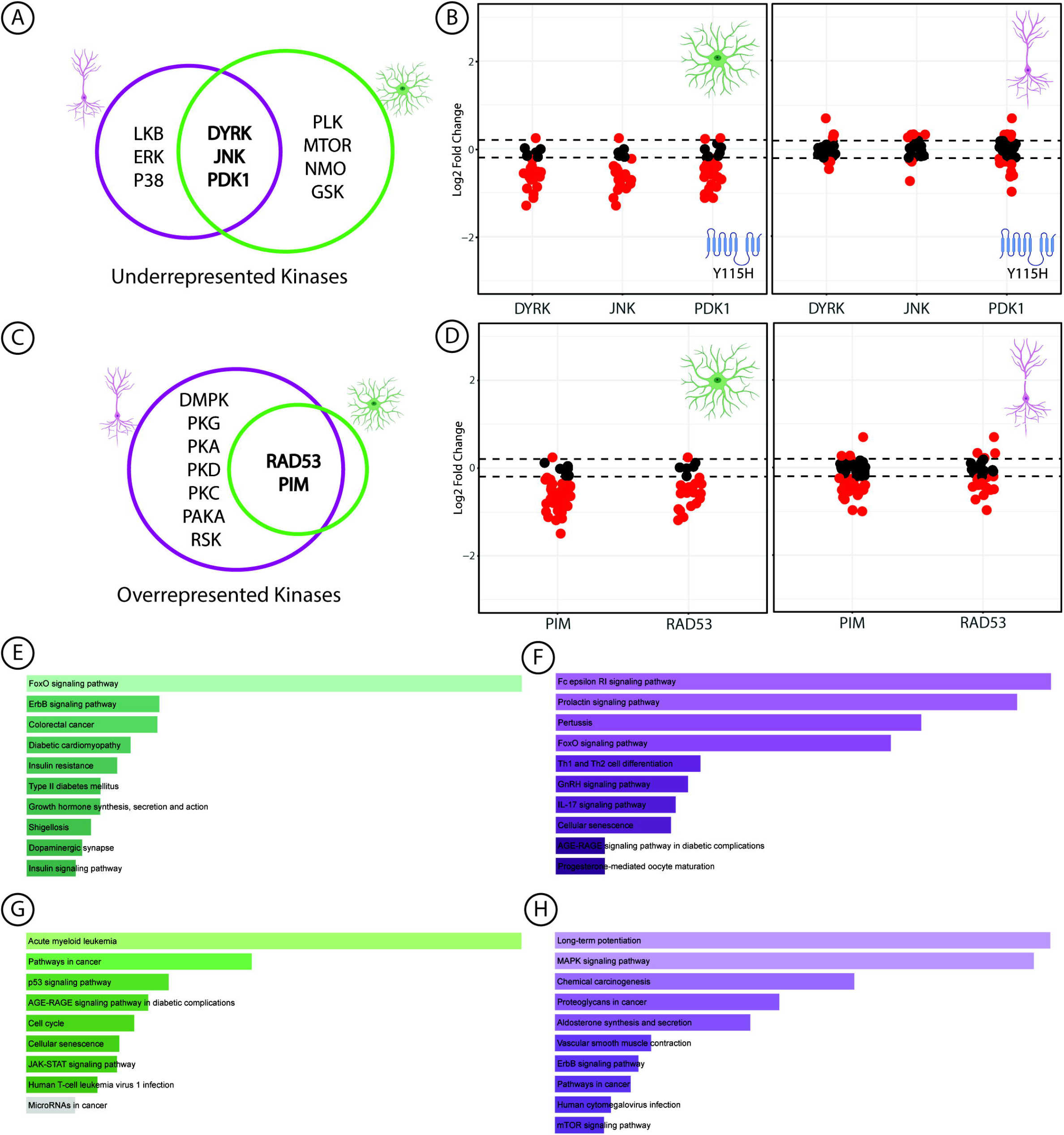
Kinases implicated in metabolic function and observed across all three familial Alzheimer’s cell lines. Our bioinformatic workflow identified both over- and underrepresented kinases that may be implicated in the etiology of AD. When analyzing all kinases with a negative Z-Score by KRSA (i.e., underrepresented kinases) across the three genotypes and our two cell types, DYRK, JNK, and PDK1 are consistently underrepresented (A). Likewise, when analyzing all kinases with a positive Z-Score by KRSA (i.e., overrepresented kinases) across the three genotypes and our cell types, RAD53 and PIM are consistently overrepresented (C). Representative peptide phosphorylation intensities of the peptides to which the three underrepresented kinases are mapped in the Male Y115H iPSCs in astrocytes (B, left) and neurons (B, right). Likewise, we have plotted the representative peptide phosphorylation intensities of the peptides to which the two overrepresented kinases are mapped in the Male Y115H iPSCs in astrocytes (D, left) and neurons (D, right). Pathway analyses of the underrepresented kinase genes in astrocytes (E) and neurons (F). and overrepresented kinase genes in astrocytes (G) and neurons (H) from Enrichr.

We extracted the genes of these five protein kinase families (Supplementary Table 2). We imported the gene names of the underrepresented and the overrepresented kinases into Enrichr. Using the KEGG 2021 Human database, we extracted the list of pathways for the common kinases across our three genotypes for both neurons and astrocytes. The most significant pathways are visualized in Figure 4E-H (Pathways are listed in Supplementary Table 3).

With the goal of identifying core-shared pathways in our substrates, we compared the top 10% significant pathways between both astrocytes and neurons for the both the under and the overrepresented kinases. For the underrepresented kinases, three pathways were shared between astrocytes and neurons: ‘FoxO signaling pathway’, ‘Prolactin signaling pathway’, and ‘Cellular senescence.’ While there was convergence on these three pathways, all of which interact downstream with canonical metabolic pathways, there were differences in the pathway enrichment. Broadly, while astrocytes were enriched in pathways for endocrine, metabolic, and human disorders of metabolism, neurons were enriched for immune function (i.e., infectious diseases and immune system).

We attempted to conduct a similar analysis for the overrepresented kinases, PIM1 and RAD53. However, due to the lack of shared kinases unique solely to astrocytes (i.e., PIM1 and RAD53 were shared overrepresented kinases between astrocytes and neurons), the analysis for astrocytes is limited to only a few genes. Interestingly, at this intersection, the most significant pathways for astrocytes and neurons related to cancer/cell growth and death pathways.

### Kinase-specific peptides phosphorylation intensities are changed for the intersection kinases

We have mapped the kinases that phosphorylate the 144 peptides on the STK chip. To determine the activity of a particular kinase, we isolated the peptides to which our intersection kinases are respectively mapped (Figure 4B, D; Supplemental Figure 3). In the astrocytes derived from the Male PSN1 Y115H cell line, 37%, 42%, and 46% of the peptides mapped to DYRK, JNK, and PDK, respectively, were under-phosphorylated while 4%, 2%, and 6% were over-phosphorylated, respectively (Figure 4B, D). In neurons derived from the Male PSN1 Y115H cell line, 20%, 20%, and 29% of the peptides mapped to DYRK, JNK, and PDK, respectively, were under-phosphorylated while 20%, 23%, and 24% were over-phosphorylated, respectively (Figure 4X). A similar analysis of the overrepresented kinases, PIM and RAD53, showed that 60% and 41% of PIM peptides were under-phosphorylated in astrocytes and neurons while 3% and 23% were over-phosphorylated, respectively. For RAD53, 64% and 52% were under-phosphorylated in astrocytes and neurons while 12% and 24% were over-phosphorylated, respectively. Kinase-specific peptide phosphorylation across all genotypes in neurons and astrocytes were perturbed (Supplemental Figure 3; Supplemental Table 4).

### In-silico and transcriptomic confirmation of the intersection kinases

To identify whether there are expression changes in our intersection kinases in postmortem Alzheimer’s Disease brain, we probed 6 large transcriptomic datasets across various brain regions for the expression of our kinase genes (Figure 5A, Supplementary Table 5). PIM1 and CHEK2 (RAD53) transcript expression was increased in the cerebellum (LFC = 0.21, p = 0.04; LFC = 0.31, p < 0.01, respectively) and the temporal cortex (LFC = 0.22, p < 0.01; LFC = 0.32, p < 0.01, respectively). CHEK2 was increased in the dlPFC (LFC = 0.13, p = 0.02, respectively). DYRK1A was increased in the cerebellum (LFC = 0.05, p = 0.04), the dlPFC (LFC = 0.04, p < 0.01), and in the parahippocampal gyrus (LFC = 0.06, p = 0.01). MAPK8 and PDK1 were decreased several across datasets. Both were decreased in the temporal cortex (LFC = −0.21, p = 0; LFC = −0.23, p < 0.01). PDK1 was decreased in the frontal pole (LFC = −0.08, p = 0.049) and while MAPK8 was decreased in the superior temporal (LFC = −0.13, p < 0.01) and the parahippocampal gyri (LFC = −0.18, p < 0.001).

**Figure 5.**
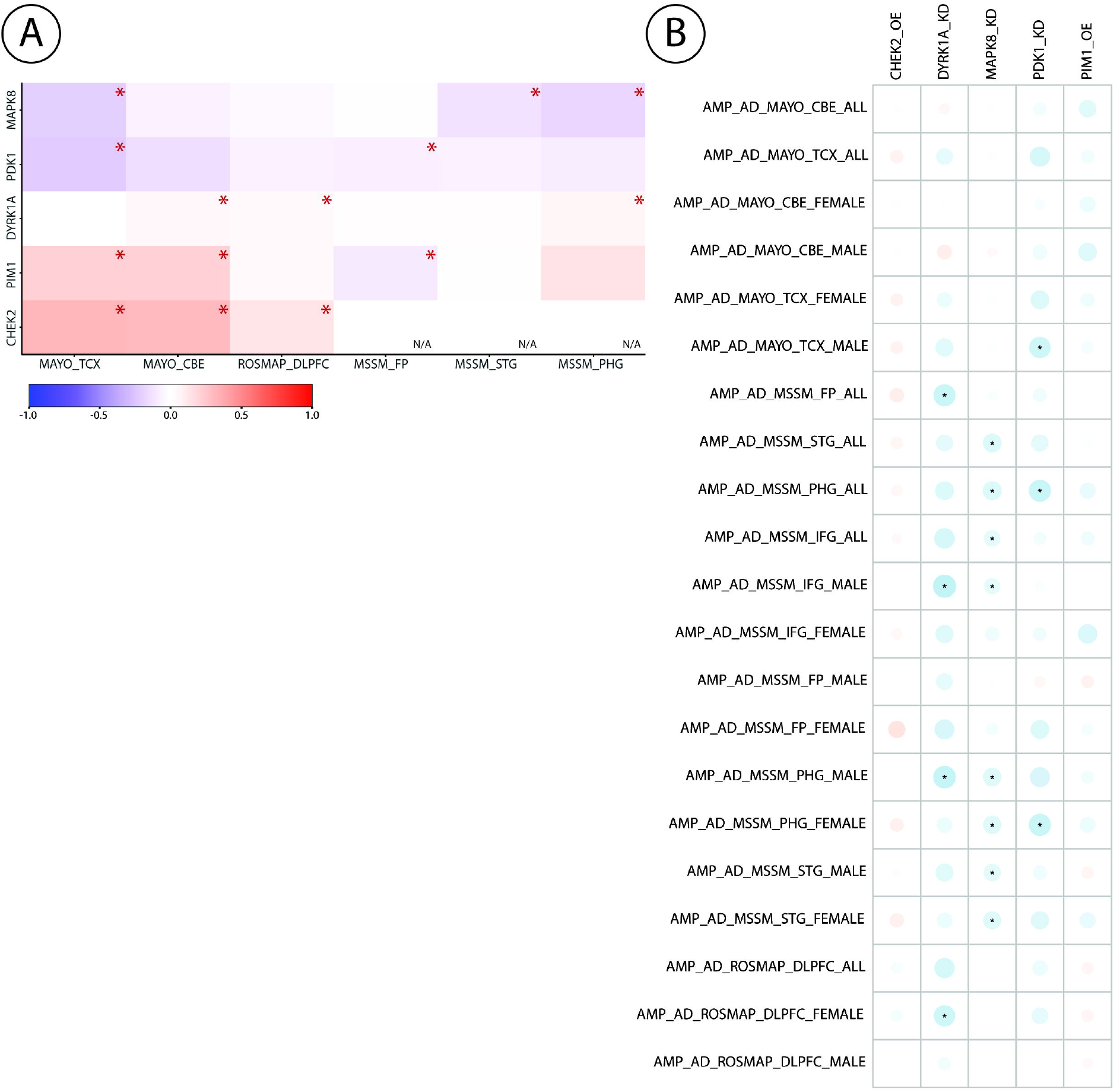
In-silico analyses of the hit AD iPSC kinases in postmortem brain. Transcript expression of the hit kinases is perturbed across various regions of the AD brain in various datasets (A). Datasets are reposited at Synapse.org (102) in the *Accelerating Medicines Partnership Program for Alzheimer’s Disease* (AMP AD): Religious Orders Study and Rush Memory and Aging Project (ROSMAP), the Mount Sinai School of Medicine (MSSM) Studies, and the Mayo Clinic (MAYO). While RAD53 (Chek2) and PIM1 are increased, PDK1 and JNK (Mapk8) are decreased consistently across brain regions (A). The red asterisk denotes significant changes in transcript expression for that respective brain region. Pearson correlation analysis of either iLINCS knockdown (MAPK8, PDK1, or DYRK1A) or overexpression (PIM1, CHEK2) signatures against 21 Alzheimer’s Disease (AD) datasets extracted from Kaleidoscope (B). The values represent correlation coefficients and color represent the sign of the coefficients (blue: positive, red: negative). The black asterisk (“*”) denotes the significance of the correlation analysis where *p* < 0.05 (B). Abbreviations are as follows: cerebellum (CBE), temporal cortex (TCX), parahippocampal gyrus (PHG), frontal pole (FP), superior temporal gyrus (STG), and dorsolateral prefrontal cortex (DLPFC).

### Connectivity analyses of intersection kinases with postmortem Alzheimer’s Disease brain

To further examine the role of our intersection kinases with the neurodegenerative disorder of Alzheimer’s disease, we performed a transcriptional connectivity analysis between gene perturbation signatures and published transcriptomic disease signatures of Alzheimer’s Disease (AD). Using the web application Kaleidoscope, we ran Pearson correlation between the iLINCS gene perturbation and AD signatures using the top differentially expressed genes in each signature (Figure 5B). Following with our KRSA assignments as well as transcriptional confirmation, we utilized three gene knockdown signatures (DYRK1A, MAPK8, and PDK1) and two overexpression signatures (PIM1 and CHEK2) in a A375 cell line. These signatures were compared to 21 AD datasets. Overall, the MAPK 8 gene signature (i.e., JNK) showed the most correlation with the AD datasets (Figure 5B; Supplemental Table 6). Eight of the 21 datasets had a significant, albeit negative, correlations in the superior temporal, parahippocampal, and inferior frontal gyri. This suggests that knockdown of MAPK8 is discordant, or opposite, the disease signature the aforementioned brain regions. That is, knocking down MAPK8 is the “reverse” of the disease and may be “therapeutic.” The overexpression signatures for our overrepresented kinases, PIM1 and RAD53 (CHEK2) had no connectivity with the AD datasets.

### DYRK1A is highly implicated in tau phosphorylation

Utilizing a previously published dataset, we assigned our intersection kinases a “tau score.” Briefly, Azorsa et al. conducted siRNA screens against the 518 protein kinases in the H4 neuroglioma cell line (40). In the knockdown conditions, both tau phosphorylation (12E8 tau) and total tau were measured. We extracted the raw data (i.e., protein expression of tau, phosphorylated tau, and the number of cells expressing the GFP-siRNA construct) and assigned “tau scores” to protein kinases. In our secondary analysis, we produce a range of tau scores. The tau score represents a normalized averaged of the phosphorylated tau/tau expression levels in the knockdown conditions. We normalized these values to have a range from 0 to 1. The lower the assigned kinase tau score, there is a higher likelihood that said kinase may play a role in the network which phosphorylates tau. The tau scores for PIM1 and RAD53 (CHEK2) was 0.33 and 0.23, respectively (Supplemental Table 7). For our underrepresented kinases, the calculated tau score for DYRK was 0.09; JNK (MAPK8), 0.19; and PDK1, 0.26. For the 518 kinases in the human genome, our calculated tau scores are listed in Supplementary Table 7. Of the 5 intersection kinases, DYRK had the lowest score (0.09) and PIM1 had the highest (0.33). Such data suggest that DYRK and its interacting network is highly likely to play a role in phosphorylating tau.

## Discussion

We identified five “core” protein kinase hits in neurons and astrocytes derived from familial AD-patient iPSCs. A clear limitation of our analyses is that we did not compare our disease cell lines to isogenic controls. Rather, we used cells isolated from non-psychiatric, non-demented patients. A future direction will be to compare the kinomes of isogenic lines to minimize stochasticity in our biological samples.

Canonically, as discussed below, these protein kinases have unique functions. We are most intrigued by the presence of kinases – at both the intersection of our comparisons and the kinases unique to either neurons or astrocytes – that are known for coordinating metabolic responses. While our pathway analysis demonstrated an enrichment of metabolic pathways specific to astrocytes, with neurons exhibiting enrichment in immune pathways, it is intriguing to consider how these intracellular networks may orchestrate metabolic signaling in AD. In the setting of cellular signals to either die or thrive, astrocytes and neurons in AD may differentially program these pathways to adapt to such changes. While our five kinases have been implicated in traditional AD pathology (such as tau phosphorylation, amyloid processing), there is emerging evidence for a role in mediating key metabolic pathways. In fact, the three shared pathways in the underrepresented kinases converge on classical pathways mediating metabolic pathways: FoxO, Prolactin, and Cellular senescence (41–51).

We summarize the evidence of our fives kinases in AD that we identified using a novel, hypothesis generating platform, completely agnostic to hypotheses that have guided the Alzheimer’s field. Importantly, our work is confirmed and complemented by a comprehensive in-silico analyses which probed postmortem RNA-Seq databases. Further research will require identifying how protein kinases such as these interact the disease progress to facilitate a neurodegenerative phenotype. As protein kinase cascades are not linear, but rather form an interconnected network, examining these hubs in such a context may provide fruitful pathophysiologic insights.

### The overrepresented kinases

#### PIM1

PIM1 is the Proviral Integration of Molony murine leukemia virus-1 (52). Largely known for its role in solid-tumor and hematologic malignancies (52), it has been studied, albeit to a lesser extent, in Alzheimer’s Disease. One study investigated the role of PIM1 in the 3xTg-AD mouse model (53). PIM1 phosphorylates PRAS40, the proline-rich AKT substrate 40 kDa, a negative regulator of the mammalian target of rapamycin (mTOR) complex (54, 55). That is, phosphorylation of PRAS40 by PIM1 disinhibits the activity of the mTOR complex. Inhibition of PIM1 improved spatial and working memory, reduced the Aβ levels, reduced tau immunoreactivity, and increased proteasome activity in this mouse model (53). Interestingly, PRAS40 is also activated by the P13K/Akt pathway (56). Because PRAS40 phosphorylation is increased in AD, and insulin signaling and hyperglycemia concomitantly facilitate its phosphorylation, the activity of PIM1 to PRAS40 may represent an evolving link in the pathophysiology of metabolic dysfunction in AD (54, 57, 58).

#### RAD53

RAD53, or CHEK2, is an important mediator of the cellular DNA damage response. CHEK2 regulates check-point entry, DNA repair, DNA replication, apoptosis, and chromatin integrity (59, 60). As a neurodegenerative disorder, examining this pathway in the pathophysiology of Alzheimer’s Disease is intriguing. This protein kinase, in the context of AD, has been largely attributed to tau hyperphosphorylation in *D. melanogaster* models (61–63). In the *Drosophila* brain, overexpression of Chek2 phosphorylated tau at Ser262, enhancing tau toxicity and tau-induced neurodegeneration (62). Likewise, flies engineered to express human amyloid-beta 42 exhibited increased tau phosphorylation at tau-Ser262 as well as increased Chek2 expression, suggesting an initiation of the DNA repair response in the presence of amyloid and tau toxicity (63). Finally, Chek2 and its counterpart Chek1 phosphorylated 27 sites on recombinant human tau (61). Of the 27 sites on tau, 13 sites have been detected in human AD brain (61). Recently, Chek1 has been implicated in AD (64). Chek1 drives the expression of an inhibitor of the protein phosphatase 2A, a major regulator of serine/threonine phosphorylation, called CIP2A (65). Both Chek1 and CIP2A protein expression is increased in the postmortem AD brain (64). Chek1 inhibition decreased tau hyperphosphorylation and inhibited amyloid production in primary neurons treated with amyloid beta oligomers (64). Further, while overexpression of Chek1 impaired working memory in the APP/PS1 mouse model, inhibition of Chek1 ameliorated cognitive function (64).

### The underrepresented kinases

#### DYRK

DYRK is the dual specificity tyrosine phosphorylation regulated kinase. Located on Chromosome 21 (66–68), it has been extensively studied in AD, especially in Down Syndrome associated AD (69, 70). DYRK1A is a key regulator of myriad neurodevelopmental processes (reviewed by (71)) and implicated in neurodegenerative disorders, tauopathies, amyloidosis, and alpha-synucleinopathies (reviewed by (72–74)). DYRK phosphorylates tau, amyloid, and presenilin (75–77). DYRK1A, was, in fact, validated in the dataset we extracted from Azorsa et al., as a “tau kinase” (40). Numerous studies have supported that DYRK inhibition either rescues, prevents, or reverses Alzheimer’s Disease pathology – both histological and behavioral – in models of the disease (78–82). Approaches to inhibit DYRK have been both genetic (81) and pharmacologic (83–87). Though there is an abundance of literature on DYRK and tau phosphorylation and amyloid pathology, the pathologic mechanism by which it functions remains unclear. There is recent evidence that DYRK undergoes aberrant proteolytic processing in the AD brain, rendering a truncated protein kinase with abnormal kinase specificity (88). These truncated forms accumulate in astrocytes and mediate a STAT3α inflammatory response (88). Additionally, DYRK modulates insulin-receptor expression in primary rat neurons (80).

#### PDK1

Pyruvate dehydrogenase complex kinase 1 (PDK1) phosphorylates and inactivates the pyruvate dehydrogenase complex, preventing the formation of Acetyl-CoA from pyruvate. Unsurprisingly, PDK1 has been studied in AD in the context of metabolic dysfunction (89–91). Vallée et al. reviewed how impaired Wnt/Beta-catenin signaling may manifest in oxidative stress and eventual cell death in neurodegenerative diseases, such as AD (89). While this pathway upregulates PDK1, in the context of AD, PDK1 expression is decreased (92). As such, Acetyl-CoA availability to the mitochondria increases, inducing oxidative stress (89). Inherently representing a theoretical mechanism by which PDK may be involved in AD, inhibition of PDK1 enhanced neuronal death in cerebellar granule neurons (93). As it relates to AD pathophysiology, PDK1 expression is decreased and postmortem frontal cortex and overexpression of PDK1 conferred neuroprotection in the B12 rat CNS cell line (90). Concordantly, in amyloid-beta resistant neuron cell lines, PDK1 expression is increased, effectively protecting the cells from toxicity (91). Finally, PDK1 transcript was differentially expressed in both a Type II Diabetes Mellitus and AD microarray dataset (94).

#### JNK

JNK (c-Jun N-terminal Kinase) is a member of the mitogen activated protein kinase (MAPK) family. There has been exhaustive research on JNK and the MAP kinases in AD (comprehensively reviewed by (95–97) and inhibitors have been designed to target these kinases (98). However, aside from the reported associations of this kinase family to canonical AD pathology, JNK has also been implicated to confer insulin resistance (99), as it phosphorylates the insulin receptor substrate 1 (100) and may impair downstream signaling through the P13K/Akt pathway (101).

## Acknowledgements

This work was supported by NIMH MH107487 and MH121102.

## Author Contributions

NDH wrote the manuscript. NDH, TM, ZW, and REM conceptualized the study. AJ, ES, and KA developed Figures from the raw kinome array data or the in-silico confirmation studies. ASI calculated the “tau scores.” AH ran the kinome array after NDH prepared the samples. CX, BX, and ZW oversaw the differentiation, growth, and maintenance of the iPSC derived cell lines. TM, ZW, and REM and REM provided critical feedback, major editorial feedback, and organized the contents of the manuscript.

---

**Supplemental Figure 1.**
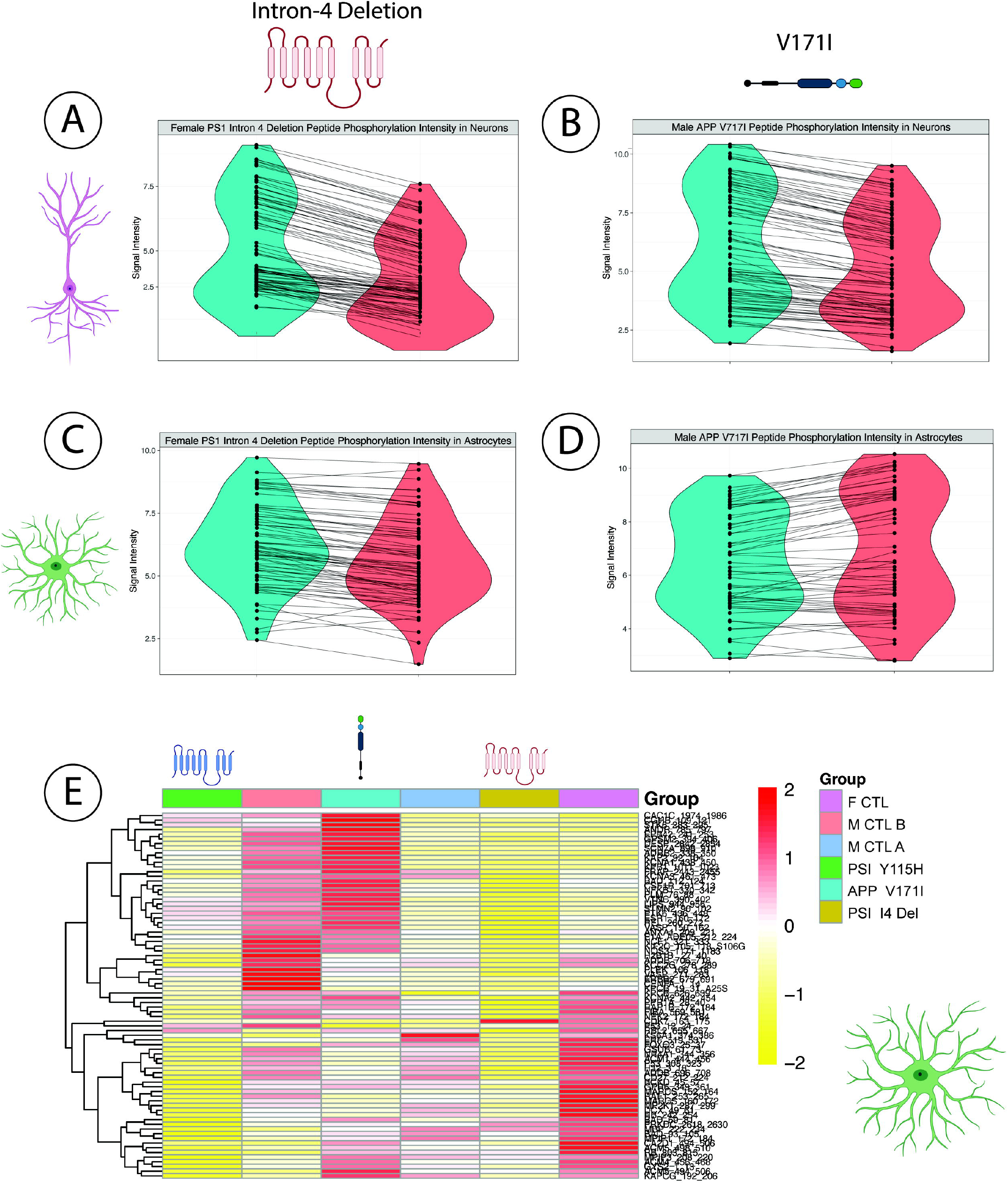
Global peptide phosphorylation is perturbed in neurons and astrocytes derived from the female PSN-1 Intron-4 deletion and the male APP V171I cell lines. In both the neurons (A) and astrocytes (C) from the female PSN-1 Intron-4 deletion subject, there is a global decrease in phosphorylation intensity. There was no difference in the global phosphorylation intensity in the neurons (B) or astrocytes (D) derived from the male V171I iPSCs. Representative heatmap of peptide phosphorylation for all astrocytes derived from our genotypes (E).

**Supplemental Figure 2.**
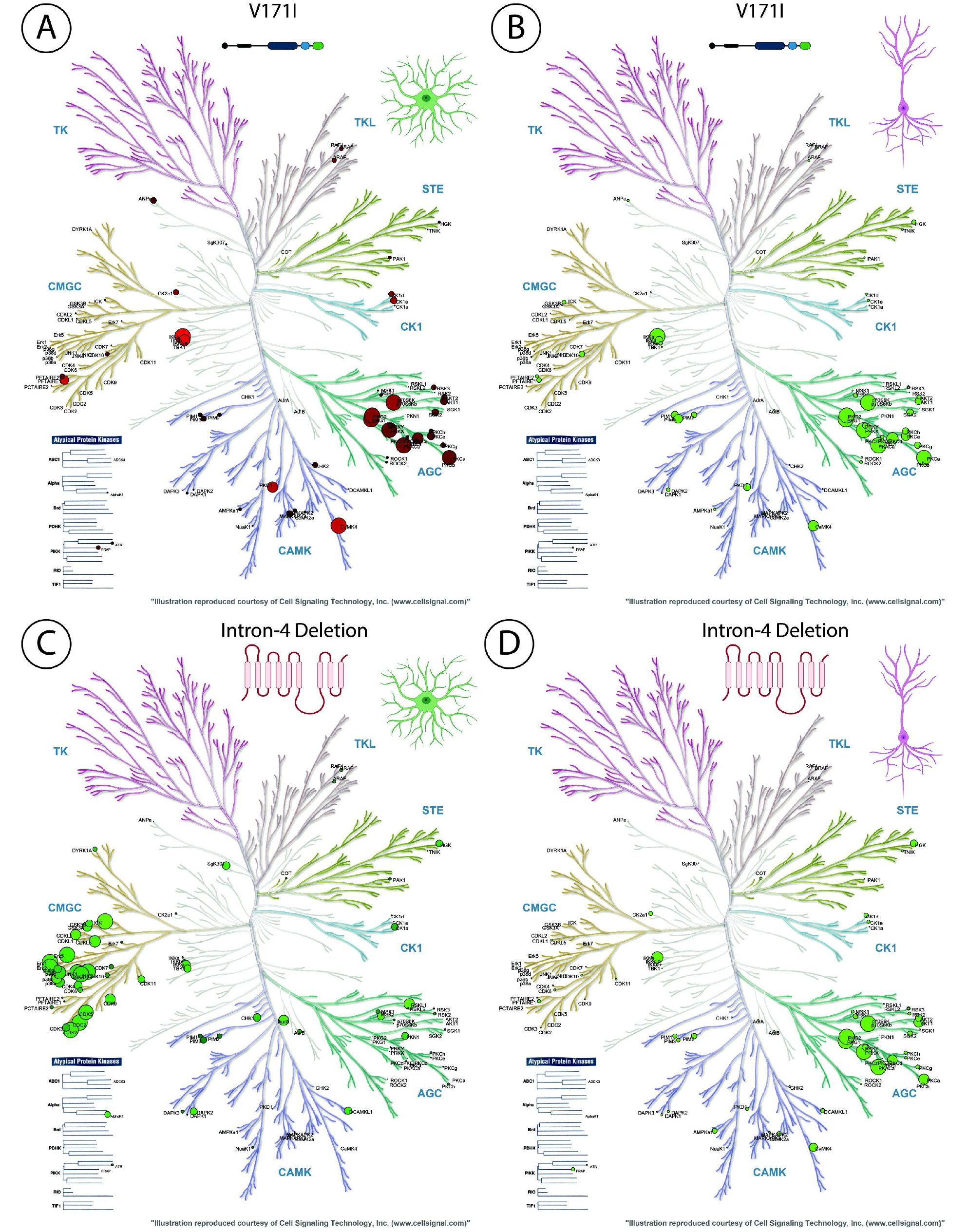
Phylogenetic Trees of the kinome in the neurons and astrocytes derived from the female PSN-1 Intron-4 deletion and the male APP V171I cell lines. While there is convergence of kinase families in the APP V171I cell line across neurons and astrocytes (A, B), the “hit” kinases are divergent in the female PSN-1 Intron-4 cell line (C, D).

**Supplemental Figure 4.**
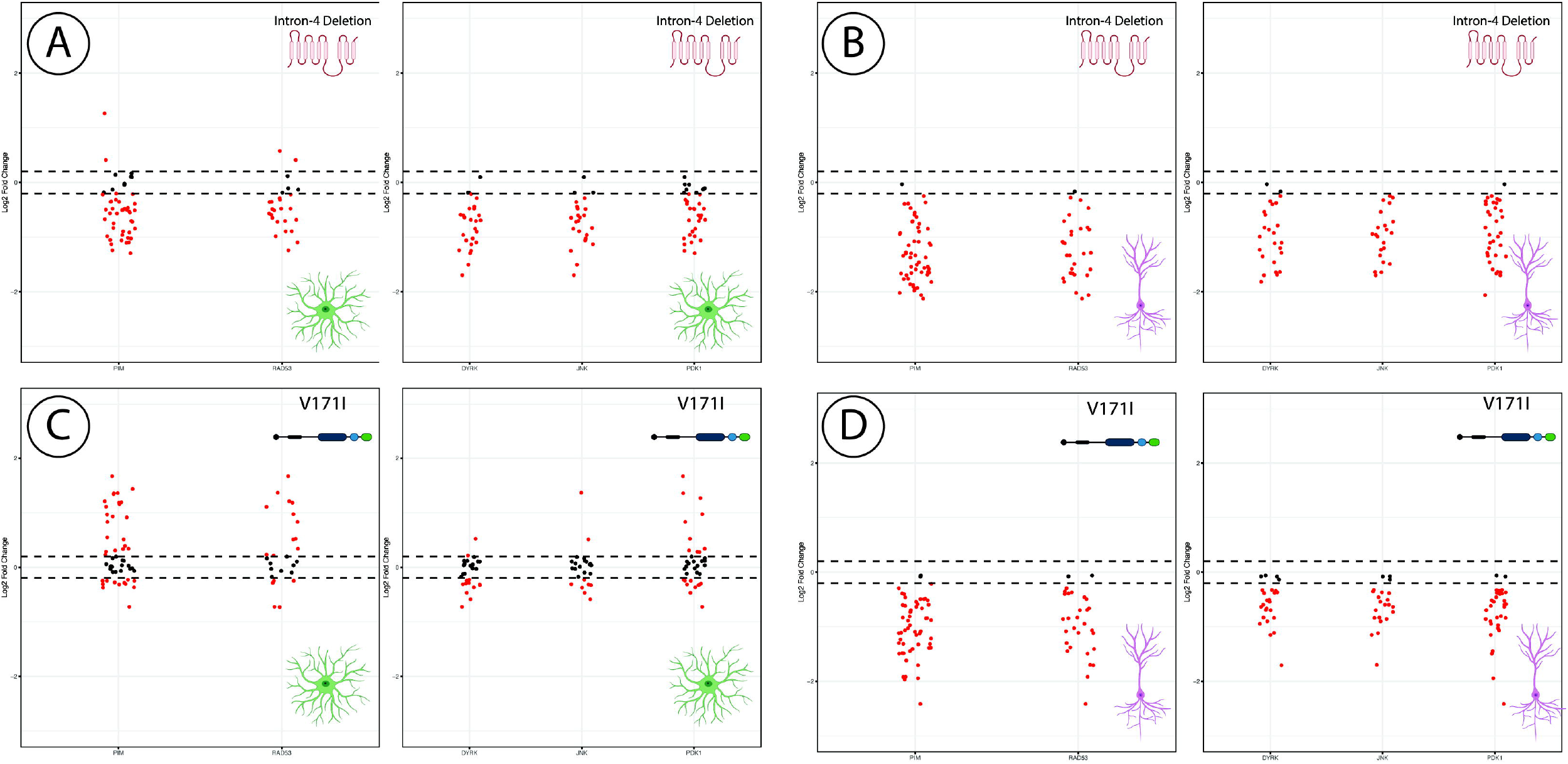
Activity of the five “hit” kinases across the neurons and astrocytes derived from the iPSCs. The Log2 Fold Change values of the peptides that are mapped to the respective kinase for the under- (left) and overrepresented (right) kinases in the female PSN1 Intron-4 deletion astrocytes (A), the female PSN1 Intron-4 deletion neurons (B), the male V717I astrocytes (C), the male V717I neurons.

## References

1. 2020 Alzheimer’s disease facts and figures. Alzheimers Dement. 2020.

2. Selkoe DJ. Cell biology of protein misfolding: the examples of Alzheimer’s and Parkinson’s diseases. Nature cell biology. 2004;6(11):1054–61.

3. Lashley T, Rohrer JD, Mead S, Revesz T. Review: an update on clinical, genetic and pathological aspects of frontotemporal lobar degenerations. Neuropathology and applied neurobiology. 2015;41(7):858–81.

4. De Strooper B, Karran E. The Cellular Phase of Alzheimer’s Disease. Cell. 2016;164(4):603–15.

5. Esparza TJ, Zhao H, Cirrito JR, Cairns NJ, Bateman RJ, Holtzman DM, et al. Amyloid-beta oligomerization in Alzheimer dementia versus high-pathology controls. Annals of neurology. 2013;73(1):104–19.

6. Jack CR, Jr., Knopman DS, Jagust WJ, Shaw LM, Aisen PS, Weiner MW, et al. Hypothetical model of dynamic biomarkers of the Alzheimer’s pathological cascade. Lancet Neurol. 2010;9(1):119–28.

7. Oksanen M, Petersen AJ, Naumenko N, Puttonen K, Lehtonen S, Gubert Olive M, et al. PSEN1 Mutant iPSC-Derived Model Reveals Severe Astrocyte Pathology in Alzheimer’s Disease. Stem Cell Reports. 2017;9(6):1885–97.

8. Ma X, Ma J, Guo R, Du J, Feng B, Liu X, et al. Blood-derived integration-free induced pluripotent stem cells (iPSCs) from one 53-years-old male donor with APOE-epsilon4/epsilon4 genotype. Stem Cell Res. 2021;54:102450.

9. Lu J, Li Y, Mollinari C, Garaci E, Merlo D, Pei G. Amyloid-beta Oligomers-induced Mitochondrial DNA Repair Impairment Contributes to Altered Human Neural Stem Cell Differentiation. Current Alzheimer research. 2019;16(10):934–49.

10. Lin YT, Seo J, Gao F, Feldman HM, Wen HL, Penney J, et al. APOE4 Causes Widespread Molecular and Cellular Alterations Associated with Alzheimer’s Disease Phenotypes in Human iPSC-Derived Brain Cell Types. Neuron. 2018;98(6):1141–54 e7.

11. Kondo T, Asai M, Tsukita K, Kutoku Y, Ohsawa Y, Sunada Y, et al. Modeling Alzheimer’s disease with iPSCs reveals stress phenotypes associated with intracellular Abeta and differential drug responsiveness. Cell stem cell. 2013;12(4):487–96.

12. Israel MA, Yuan SH, Bardy C, Reyna SM, Mu Y, Herrera C, et al. Probing sporadic and familial Alzheimer’s disease using induced pluripotent stem cells. Nature. 2012;482(7384):216–20.

13. Elsworthy RJ, King MC, Grainger A, Fisher E, Crowe JA, Alqattan S, et al. Amyloid-beta precursor protein processing and oxidative stress are altered in human iPSC-derived neuron and astrocyte co-cultures carrying presenillin-1 gene mutations following spontaneous differentiation. Molecular and cellular neurosciences. 2021;114:103631.

14. Dubey SK, Ram MS, Krishna KV, Saha RN, Singhvi G, Agrawal M, et al. Recent Expansions on Cellular Models to Uncover the Scientific Barriers Towards Drug Development for Alzheimer’s Disease. Cellular and molecular neurobiology. 2019;39(2):181–209.

15. Barak M, Fedorova V, Pospisilova V, Raska J, Vochyanova S, Sedmik J, et al. Human iPSC-Derived Neural Models for Studying Alzheimer’s Disease: from Neural Stem Cells to Cerebral Organoids. Stem Cell Rev Rep. 2022;18(2):792–820.

16. Albert K, Niskanen J, Kalvala S, Lehtonen S. Utilising Induced Pluripotent Stem Cells in Neurodegenerative Disease Research: Focus on Glia. Int J Mol Sci. 2021;22(9).

17. Huang LK, Chao SP, Hu CJ. Clinical trials of new drugs for Alzheimer disease. J Biomed Sci. 2020;27(1):18.

18. Terry M. A Long Line of Alzheimer’s Failures: Roche Drops Two Drug Trials: BioSpace; 2019 [Available from: https://www.biospace.com/article/a-long-line-of-failures-roche-drops-alzheimer-s-drug-trials/.

19. Alganem K, Hamoud AR, Creeden JF, Henkel ND, Imami AS, Joyce AW, et al. The active kinome: The modern view of how active protein kinase networks fit in biological research. Curr Opin Pharmacol. 2022;62:117–29.

20. Wang C, Yu JT, Miao D, Wu ZC, Tan MS, Tan L. Targeting the mTOR signaling network for Alzheimer’s disease therapy. Mol Neurobiol. 2014;49(1):120–35.

21. Caberlotto L, Lauria M, Nguyen TP, Scotti M. The central role of AMP-kinase and energy homeostasis impairment in Alzheimer’s disease: a multifactor network analysis. PloS one. 2013;8(11):e78919.

22. Rosenberger AF, Hilhorst R, Coart E, Garcia Barrado L, Naji F, Rozemuller AJ, et al. Protein Kinase Activity Decreases with Higher Braak Stages of Alzheimer’s Disease Pathology. Journal of Alzheimer’s disease: JAD. 2016;49(4):927–43.

23. Lanke V, Moolamalla STR, Roy D, Vinod PK. Integrative Analysis of Hippocampus Gene Expression Profiles Identifies Network Alterations in Aging and Alzheimer’s Disease. Front Aging Neurosci. 2018;10:153.

24. Liang D, Han G, Feng X, Sun J, Duan Y, Lei H. Concerted perturbation observed in a hub network in Alzheimer’s disease. PloS one. 2012;7(7):e40498.

25. Hallock P, Thomas MA. Integrating the Alzheimer’s disease proteome and transcriptome: a comprehensive network model of a complex disease. OMICS. 2012;16(1-2):37–49.

26. Petyuk VA, Chang R, Ramirez-Restrepo M, Beckmann ND, Henrion MYR, Piehowski PD, et al. The human brainome: network analysis identifies HSPA2 as a novel Alzheimer’s disease target. Brain: a journal of neurology. 2018;141(9):2721–39.

27. Bai B, Wang X, Li Y, Chen PC, Yu K, Dey KK, et al. Deep Multilayer Brain Proteomics Identifies Molecular Networks in Alzheimer’s Disease Progression. Neuron. 2020;105(6):975–91 e7.

28. Johnson ECB, Carter EK, Dammer EB, Duong DM, Gerasimov ES, Liu Y, et al. Large-scale deep multi-layer analysis of Alzheimer’s disease brain reveals strong proteomic disease-related changes not observed at the RNA level. Nat Neurosci. 2022;25(2):213–25.

29. Wen Z, Nguyen HN, Guo Z, Lalli MA, Wang X, Su Y, et al. Synaptic dysregulation in a human iPS cell model of mental disorders. Nature. 2014;515(7527):414–8.

30. Xu M, Lee EM, Wen Z, Cheng Y, Huang WK, Qian X, et al. Identification of small-molecule inhibitors of Zika virus infection and induced neural cell death via a drug repurposing screen. Nature medicine. 2016;22(10):1101–7.

31. Kim NS, Wen Z, Liu J, Zhou Y, Guo Z, Xu C, et al. Pharmacological rescue in patient iPSC and mouse models with a rare DISC1 mutation. Nature communications. 2021;12(1):1398.

32. Bentea E, Depasquale EAK, O’Donovan SM, Sullivan CR, Simmons M, Meador-Woodruff JH, et al. Kinase network dysregulation in a human induced pluripotent stem cell model of DISC1 schizophrenia. Mol Omics. 2019;15(3):173–88.

33. DePasquale EAK, Alganem K, Bentea E, Nawreen N, McGuire JL, Tomar T, et al. KRSA: An R package and R Shiny web application for an end-to-end upstream kinase analysis of kinome array data. PloS one. 2021;16(12):e0260440.

34. Chen EY, Tan CM, Kou Y, Duan Q, Wang Z, Meirelles GV, et al. Enrichr: interactive and collaborative HTML5 gene list enrichment analysis tool. BMC bioinformatics. 2013;14:128.

35. Genomics VUBE. [Available from: https://bioinformatics.psb.ugent.be/webtools/Venn/.

36. Asah S, Alganem K, McCullumsmith RE, O’Donovan SM. A bioinformatic inquiry of the EAAT2 interactome in postmortem and neuropsychiatric datasets. Schizophrenia research. 2020.

37. [Available from: https://kalganem.shinyapps.io/BrainDatabases/.

38. Pilarczyk M, Kouril M, Shamsaei B, Vasiliauskas J, Niu W, Mahi N, et al. Connecting omics signatures of diseases, drugs, and mechanisms of actions with iLINCS. bioRxiv. 2020:826271.

39. KinMap [Available from: http://www.kinhub.org/kinmap/.

40. Azorsa DO, Robeson RH, Frost D, Meec hoovet B, Brautigam GR, Dickey C, et al. High-content siRNA screening of the kinome identifies kinases involved in Alzheimer’s disease-related tau hyperphosphorylation. BMC genomics. 2010;11:25.

41. von Zglinicki T, Wan T, Miwa S. Senescence in Post-Mitotic Cells: A Driver of Aging? Antioxidants & redox signaling. 2021;34(4):308–23.

42. Triebel J, Robles JP, Zamora M, Clapp C, Bertsch T. New horizons in specific hormone proteolysis. Trends Endocrinol Metab. 2022;33(6):371–7.

43. Sun EJ, Wankell M, Palamuthusingam P, McFarlane C, Hebbard L. Targeting the PI3K/Akt/mTOR Pathway in Hepatocellular Carcinoma. Biomedicines. 2021;9(11).

44. Rapaka D, Bitra VR, Challa SR, Adiukwu PC. mTOR signaling as a molecular target for the alleviation of Alzheimer’s disease pathogenesis. Neurochemistry international. 2022;155:105311.

45. Polsky LR, Rentscher KE, Carroll JE. Stress-induced biological aging: A review and guide for research priorities. Brain, behavior, and immunity. 2022;104:97–109.

46. Molina-Salinas G, Rivero-Segura NA, Cabrera-Reyes EA, Rodriguez-Chavez V, Langley E, Cerbon M. Decoding signaling pathways involved in prolactin-induced neuroprotection: A review. Front Neuroendocrinol. 2021;61:100913.

47. Maiese K. Neurodegeneration, memory loss, and dementia: the impact of biological clocks and circadian rhythm. Front Biosci (Landmark Ed). 2021;26(9):614–27.

48. Ge Y, Zhou M, Chen C, Wu X, Wang X. Role of AMPK mediated pathways in autophagy and aging. Biochimie. 2022;195:100–13.

49. Gass S, Harris J, Ormandy C, Brisken C. Using gene expression arrays to elucidate transcriptional profiles underlying prolactin function. J Mammary Gland Biol Neoplasia. 2003;8(3):269–85.

50. Du S, Zheng H. Role of FoxO transcription factors in aging and age-related metabolic and neurodegenerative diseases. Cell Biosci. 2021;11(1):188.

51. Chao CC, Shen PW, Tzeng TY, Kung HJ, Tsai TF, Wong YH. Human iPSC-Derived Neurons as A Platform for Deciphering the Mechanisms behind Brain Aging. Biomedicines. 2021;9(11).

52. Zhao Y, Aziz AUR, Zhang H, Zhang Z, Li N, Liu B. A systematic review on active sites and functions of PIM-1 protein. Hum Cell. 2022;35(2):427–40.

53. Velazquez R, Shaw DM, Caccamo A, Oddo S. Pim1 inhibition as a novel therapeutic strategy for Alzheimer’s disease. Mol Neurodegener. 2016;11(1):52.

54. Sancak Y, Thoreen CC, Peterson TR, Lindquist RA, Kang SA, Spooner E, et al. PRAS40 is an insulin-regulated inhibitor of the mTORC1 protein kinase. Molecular cell. 2007;25(6):903–15.

55. Wang L, Harris TE, Roth RA, Lawrence JC, Jr. PRAS40 regulates mTORC1 kinase activity by functioning as a direct inhibitor of substrate binding. The Journal of biological chemistry. 2007;282(27):20036–44.

56. Wang YF, Khan M, van den Berg HA. Interaction of fast and slow dynamics in endocrine control systems with an application to beta-cell dynamics. Math Biosci. 2012;235(1):8–18.

57. Ma Y, Wu D, Zhang W, Liu J, Chen S, Hua B. Investigation of PI3K/PKB/mTOR/S6K1 signaling pathway in relationship of type 2 diabetes and Alzheimer’s disease. Int J Clin Exp Med. 2015;8(10):18581–90.

58. Nascimento EB, Fodor M, van der Zon GC, Jazet IM, Meinders AE, Voshol PJ, et al. Insulin-mediated phosphorylation of the proline-rich Akt substrate PRAS40 is impaired in insulin target tissues of high-fat diet-fed rats. Diabetes. 2006;55(12):3221–8.

59. Reinhardt HC, Yaffe MB. Kinases that control the cell cycle in response to DNA damage: Chk1, Chk2, and MK2. Curr Opin Cell Biol. 2009;21(2):245–55.

60. Stracker TH, Usui T, Petrini JH. Taking the time to make important decisions: the checkpoint effector kinases Chk1 and Chk2 and the DNA damage response. DNA Repair (Amst). 2009;8(9):1047–54.

61. Mendoza J, Sekiya M, Taniguchi T, Iijima KM, Wang R, Ando K. Global analysis of phosphorylation of tau by the checkpoint kinases Chk1 and Chk2 in vitro. Journal of proteome research. 2013;12(6):2654–65.

62. Iijima-Ando K, Zhao L, Gatt A, Shenton C, Iijima K. A DNA damage-activated checkpoint kinase phosphorylates tau and enhances tau-induced neurodegeneration. Human molecular genetics. 2010;19(10):1930–8.

63. Iijima K, Gatt A, Iijima-Ando K. Tau Ser262 phosphorylation is critical for Abeta42-induced tau toxicity in a transgenic Drosophila model of Alzheimer’s disease. Human molecular genetics. 2010;19(15):2947–57.

64. Hu W, Wang Z, Zhang H, Mahaman YAR, Huang F, Meng D, et al. Chk1 Inhibition Ameliorates Alzheimer’s Disease Pathogenesis and Cognitive Dysfunction Through CIP2A/PP2A Signaling. Neurotherapeutics: the journal of the American Society for Experimental NeuroTherapeutics. 2022.

65. Khanna A, Kauko O, Bockelman C, Laine A, Schreck I, Partanen JI, et al. Chk1 targeting reactivates PP2A tumor suppressor activity in cancer cells. Cancer research. 2013;73(22):6757–69.

66. Shindoh N, Kudoh J, Maeda H, Yamaki A, Minoshima S, Shimizu Y, et al. Cloning of a human homolog of the Drosophila minibrain/rat Dyrk gene from “the Down syndrome critical region” of chromosome 21. Biochemical and biophysical research communications. 1996;225(1):92–9.

67. Kentrup H, Becker W, Heukelbach J, Wilmes A, Schurmann A, Huppertz C, et al. Dyrk, a dual specificity protein kinase with unique structural features whose activity is dependent on tyrosine residues between subdomains VII and VIII. The Journal of biological chemistry. 1996;271(7):3488–95.

68. Guimera J, Casas C, Pucharcos C, Solans A, Domenech A, Planas AM, et al. A human homologue of Drosophila minibrain (MNB) is expressed in the neuronal regions affected in Down syndrome and maps to the critical region. Human molecular genetics. 1996;5(9):1305–10.

69. Park J, Song WJ, Chung KC. Function and regulation of Dyrk1A: towards understanding Down syndrome. Cell Mol Life Sci. 2009;66(20):3235–40.

70. Kimura R, Kamino K, Yamamoto M, Nuripa A, Kida T, Kazui H, et al. The DYRK1A gene, encoded in chromosome 21 Down syndrome critical region, bridges between beta-amyloid production and tau phosphorylation in Alzheimer disease. Human molecular genetics. 2007;16(1):15–23.

71. Tejedor FJ, Hammerle B. MNB/DYRK1A as a multiple regulator of neuronal development. The FEBS journal. 2011;278(2):223–35.

72. Wegiel J, Gong CX, Hwang YW. The role of DYRK1A in neurodegenerative diseases. The FEBS journal. 2011;278(2):236–45.

73. Pathak A, Rohilla A, Gupta T, Akhtar MJ, Haider MR, Sharma K, et al. DYRK1A kinase inhibition with emphasis on neurodegeneration: A comprehensive evolution story-cum-perspective. Eur J Med Chem. 2018;158:559–92.

74. Ferrer I, Barrachina M, Puig B, Martinez de Lagran M, Marti E, Avila J, et al. Constitutive Dyrk1A is abnormally expressed in Alzheimer disease, Down syndrome, Pick disease, and related transgenic models. Neurobiology of disease. 2005;20(2):392–400.

75. Ryoo SR, Jeong HK, Radnaabazar C, Yoo JJ, Cho HJ, Lee HW, et al. DYRK1A-mediated hyperphosphorylation of Tau. A functional link between Down syndrome and Alzheimer disease. The Journal of biological chemistry. 2007;282(48):34850–7.

76. Ryu YS, Park SY, Jung MS, Yoon SH, Kwen MY, Lee SY, et al. Dyrk1A-mediated phosphorylation of Presenilin 1: a functional link between Down syndrome and Alzheimer’s disease. Journal of neurochemistry. 2010;115(3):574–84.

77. Liu F, Liang Z, Wegiel J, Hwang YW, Iqbal K, Grundke-Iqbal I, et al. Overexpression of Dyrk1A contributes to neurofibrillary degeneration in Down syndrome. FASEB journal: official publication of the Federation of American Societies for Experimental Biology. 2008;22(9):3224–33.

78. Velazquez R, Meechoovet B, Ow A, Foley C, Shaw A, Smith B, et al. Chronic Dyrk1 Inhibition Delays the Onset of AD-Like Pathology in 3xTg-AD Mice. Mol Neurobiol. 2019;56(12):8364–75.

79. Stotani S, Giordanetto F, Medda F. DYRK1A inhibition as potential treatment for Alzheimer’s disease. Future Med Chem. 2016;8(6):681–96.

80. Souchet B, Audrain M, Billard JM, Dairou J, Fol R, Orefice NS, et al. Inhibition of DYRK1A proteolysis modifies its kinase specificity and rescues Alzheimer phenotype in APP/PS1 mice. Acta Neuropathol Commun. 2019;7(1):46.

81. Liu Y, Wang L, Xie F, Wang X, Hou Y, Wang X, et al. Overexpression of miR-26a-5p Suppresses Tau Phosphorylation and Abeta Accumulation in the Alzheimer’s Disease Mice by Targeting DYRK1A. Curr Neurovasc Res. 2020;17(3):241–8.

82. Branca C, Shaw DM, Belfiore R, Gokhale V, Shaw AY, Foley C, et al. Dyrk1 inhibition improves Alzheimer’s disease-like pathology. Aging Cell. 2017;16(5):1146–54.

83. Smith B, Medda F, Gokhale V, Dunckley T, Hulme C. Recent advances in the design, synthesis, and biological evaluation of selective DYRK1A inhibitors: a new avenue for a disease modifying treatment of Alzheimer’s? ACS Chem Neurosci. 2012;3(11):857–72.

84. Naert G, Ferre V, Meunier J, Keller E, Malmstrom S, Givalois L, et al. Leucettine L41, a DYRK1A-preferential DYRKs/CLKs inhibitor, prevents memory impairments and neurotoxicity induced by oligomeric Abeta25-35 peptide administration in mice. European neuropsychopharmacology: the journal of the European College of Neuropsychopharmacology. 2015;25(11):2170–82.

85. Demuro S, Sauvey C, Tripathi SK, Di Martino RMC, Shi D, Ortega JA, et al. ARN25068, a versatile starting point towards triple GSK-3beta/FYN/DYRK1A inhibitors to tackle tau-related neurological disorders. Eur J Med Chem. 2022;229:114054.

86. Coutadeur S, Benyamine H, Delalonde L, de Oliveira C, Leblond B, Foucourt A, et al. A novel DYRK1A (dual specificity tyrosine phosphorylation-regulated kinase 1A) inhibitor for the treatment of Alzheimer’s disease: effect on Tau and amyloid pathologies in vitro. Journal of neurochemistry. 2015;133(3):440–51.

87. Lee HJ, Woo H, Lee HE, Jeon H, Ryu KY, Nam JH, et al. The novel DYRK1A inhibitor KVN93 regulates cognitive function, amyloid-beta pathology, and neuroinflammation. Free radical biology & medicine. 2020;160:575–95.

88. Tian S, Jia W, Lu M, Zhao J, Sun X. Dual-specificity tyrosine phosphorylation-regulated kinase 1A ameliorates insulin resistance in neurons by up-regulating IRS-1 expression. The Journal of biological chemistry. 2019;294(52):20164–76.

89. Vallee A, Lecarpentier Y, Guillevin R, Vallee JN. Thermodynamics in Neurodegenerative Diseases: Interplay Between Canonical WNT/Beta-Catenin Pathway-PPAR Gamma, Energy Metabolism and Circadian Rhythms. Neuromolecular Med. 2018;20(2):174–204.

90. Newington JT, Rappon T, Albers S, Wong DY, Rylett RJ, Cumming RC. Overexpression of pyruvate dehydrogenase kinase 1 and lactate dehydrogenase A in nerve cells confers resistance to amyloid beta and other toxins by decreasing mitochondrial respiration and reactive oxygen species production. The Journal of biological chemistry. 2012;287(44):37245–58.

91. Newington JT, Pitts A, Chien A, Arseneault R, Schubert D, Cumming RC. Amyloid beta resistance in nerve cell lines is mediated by the Warburg effect. PloS one. 2011;6(4):e19191.

92. Pate KT, Stringari C, Sprowl-Tanio S, Wang K, TeSlaa T, Hoverter NP, et al. Wnt signaling directs a metabolic program of glycolysis and angiogenesis in colon cancer. EMBO J. 2014;33(13):1454–73.

93. Bobba A, Amadoro G, La Piana G, Calissano P, Atlante A. Glycolytic enzyme upregulation and numbness of mitochondrial activity characterize the early phase of apoptosis in cerebellar granule cells. Apoptosis: an international journal on programmed cell death. 2015;20(1):10–28.

94. Mirza Z, Kamal MA, Buzenadah AM, Al-Qahtani MH, Karim S. Establishing genomic/transcriptomic links between Alzheimer’s disease and type 2 diabetes mellitus by meta-analysis approach. CNS Neurol Disord Drug Targets. 2014;13(3):501–16.

95. Hugon J, Mouton-Liger F, Cognat E, Dumurgier J, Paquet C. Blood-Based Kinase Assessments in Alzheimer’s Disease. Front Aging Neurosci. 2018;10:338.

96. Yarza R, Vela S, Solas M, Ramirez MJ. c-Jun N-terminal Kinase (JNK) Signaling as a Therapeutic Target for Alzheimer’s Disease. Front Pharmacol. 2015;6:321.

97. Ahmed T, Zulfiqar A, Arguelles S, Rasekhian M, Nabavi SF, Silva AS, et al. Map kinase signaling as therapeutic target for neurodegeneration. Pharmacological research: the official journal of the Italian Pharmacological Society. 2020;160:105090.

98. Li G, Qi W, Li X, Zhao J, Luo M, Chen J. Recent Advances in c-Jun N-Terminal Kinase (JNK) Inhibitors. Curr Med Chem. 2021;28(3):607–27.

99. Wang Q, Duan L, Li X, Wang Y, Guo W, Guan F, et al. Glucose Metabolism, Neural Cell Senescence and Alzheimer’s Disease. Int J Mol Sci. 2022;23(8).

100. Sabio G, Das M, Mora A, Zhang Z, Jun JY, Ko HJ, et al. A stress signaling pathway in adipose tissue regulates hepatic insulin resistance. Science. 2008;322(5907):1539–43.

101. Bomfim TR, Forny-Germano L, Sathler LB, Brito-Moreira J, Houzel JC, Decker H, et al. An anti-diabetes agent protects the mouse brain from defective insulin signaling caused by Alzheimer’s disease-associated Abeta oligomers. The Journal of clinical investigation. 2012;122(4):1339–53.

102. AMP-AD: Accelerating Medicines Partnership - Alzheimer’s Disease Target Discovery and Preclinical Validation [Available from: https://adknowledgeportal.synapse.org/Explore/Programs/DetailsPage?Program=AMP-AD.

